# MacaquePose: A novel ‘in the wild’ macaque monkey pose dataset for markerless motion capture

**DOI:** 10.1101/2020.07.30.229989

**Authors:** Rollyn Labuguen, Jumpei Matsumoto, Salvador Negrete, Hiroshi Nishimaru, Hisao Nishijo, Masahiko Takada, Yasuhiro Go, Ken-ichi Inoue, Tomohiro Shibata

## Abstract

Video-based markerless motion capture permits quantification of an animal’s pose and motion, with a high spatiotemporal resolution in a naturalistic context, and is a powerful tool for analyzing the relationship between the animal’s behaviors and its brain functions. Macaque monkeys are excellent non-human primate models, especially for studying neuroscience. Due to the lack of a dataset allowing training of a deep neural network for the macaque’s markerless motion capture in the naturalistic context, it has been challenging to apply this technology for macaques-based studies. In this study, we created MacaquePose, a novel open dataset with manually labeled body part positions for macaques in naturalistic scenes, consisting of >13,000 images, refined by researchers. We show that the pose estimation performance of an artificial neural network trained with the dataset is close to that of a human-level. The MacaquePose will provide a platform for innovative behavior analysis for non-human primate.

## 1 Introduction

Behavior analyses are fundamental for understanding brain functions and malfunctions (Datta et al., 2019). Motion capture technologies allow the quantification of animal’s pose and motion with a high spatiotemporal resolution enabling the study of the relationship between various brain functions and behaviors (Vargas-Irwin et al., 2008; Nagasaka et al., 2011; Mathis et al., 2020). However, attaching the physical markers for the motion capture is often not practical for animal studies, as the markers themselves disturb/change the subject’s behavior (Nakamura et al., 2016; Mathis et al., 2018; Berger et al., 2020). Thanks to recent advances in machine vision using deep learning, the video-based markerless motion capture has been developed to a level permitting practical use (Mathis et al., 2020), in which an artificial neural network predicts the location of body parts in a video without the requirement for physical markers, and enabled successful behavioral studies in rodents (e.g., Dooley et al., 2020; Cregg et al., 2020; Mathis et al., 2020). Macaque monkeys are an important non-human primate model, particularly in the field of neuroscience (Kalin et al., 2006; Capitanio et al., 2008; Nelson et al., 2008; Watson et al., 2012). The robust markerless motion capture using deep learning will allow studying various complex naturalistic behaviors in detail, and permit investigation of relationship between naturalistic behaviors and brain functions (Datta et al., 2019; Mathis et al., 2020). Analyzing naturalistic behavior is crucial in brain-science, since the brain evolved from natural behaviors, and various behaviors, such as complex social behaviors, can be observed only in the natural situations (Datta et al., 2019; Mathis et al., 2020). The deep neural networks usually require manually labeled body parts positions in thousands of pictures to learn prediction of the body parts positions in an arbitrary picture. However, such a large labeled dataset for macaque monkeys in the naturalistic scene has not been developed. The lack of this dataset limits the markerless motion capture technology applications for macaque studies (Berger et al., 2020; Bala et al., 2020)

To overcome this limitation, we created a novel open dataset of the manually labeled body part positions (keypoints) for macaques in naturalistic scenes, consisting of greater than 13,000 pictures. We also validated the usefulness of the dataset by training and evaluating an artificial neural network with the dataset. The results revealed that the keypoint estimation performance of the trained network was close to that of a human level. Our dataset will provide basis for markerless motion capture on the naturalistic behaviors.

## 2 Materials and Methods

### 2.1 Image Data Collection

A total of 13,083 images of macaque monkeys were obtained from the internet or were captured in zoos or the Primate Research Institute of Kyoto University. Images on the internet were obtained through Google Open Images (https://storage.googleapis.com/openimages/web/index.html) by searching for images with a ‘macaque’ tag. Pictures zoos were acquired from the outside of the breeding areas, with granted permission provided by the zoos. Images in the Primate Research Institute of Kyoto University were taken in the breeding fields without causing any specific interventions to the monkeys. The photo capturing in the institute was approved by the Animal Welfare and Animal Care Committee of the Primate Research Institute of Kyoto University and conducted in accordance with the Guidelines for the Care and Use of Animals of the Primate Research Institute, Kyoto University.

### 2.2 Image Data Annotation

The positions of 17 keypoints (nose and left and right ears, eyes, shoulders, elbows, wrists, hips, knees, and ankles) and instance segmentation for each monkey in each of the pictures were first annotated by non-researchers employed by Baobab Inc. (Chiyoda-ku, Japan). As further expertise was required for high-quality monkey annotation, the keypoint labels were then further refined with eight researchers working with macaques at Kyoto University and the University of Toyama, using a custom-made Python script. The keypoints were labeled according to the following guidelines: 1) The keypoints of the limbs (shoulder, elbow, wrist, hip, knee, and ankle) should be located at the center of the joint rotation. 2) Ear, eye, and nose keypoints should be located at the entrance of the ear canal, the center of eye ball, in the middle position between the entrances of the two nostrils, respectively. 3) A keypoint was annotated, if its position was predictable despite being occluded, except for ears, eyes, and nose facing the back side of the picture. The resultant labels were compatible with the Microsoft COCO Keypoint Dataset (Lin et al., 2014).

### 2.3 Performance evaluation of an artificial neural network trained with the present dataset

To validate the present dataset, we trained an artificial neural network estimating keypoint positions by using the DeepLabCut algorithm (Mathis et al., 2018). Briefly, DeepLabCut is a versatile and straightforward algorithm in which the 50-layer ResNet pre-trained for the ImageNet object recognition task (He et al., 2016) is transferred for the keypoint estimation by replacing the classification layer at the output of the ResNet with the deconvolutional layers. DeepLabCut is a widely used algorithm in the field of neuroscience as it requires a relatively small number of training data, has user-friendly interface associated with the algorithm, and has a proven good performance (Nath et al., 2019). Due to DeepLabCut (version 2.1.6) currently not supporting the estimation of keypoints in multiple animals in a picture, we first generated single monkey images by masking the monkeys in the images except for one monkey and used these masked images as the input. Some monkey images in the dataset were excluded due to technical reasons (e.g., a keypoint of one monkey is covered by the mask of the other monkeys). Then, the images were resized to adjust the length to 640 pixels while maintaining the images aspect ratio, before inputting it into the network. In total, 15,476 single monkey images were generated. Among the images, 14,697 single monkey images were used to train the network and the rest (779 images) were used to evaluate the trained network.

The network is trained up to a million iterations. The training took 20 hours to complete on a Nvidia GTX 1080 Ti graphics processing unit workstation. The keypoint prediction by the trained network was evaluated. A predicted keypoint with confidence level > 0.4 was defined to be detected. First, minor cases showing the keypoint(s) detected outside the monkey segment were eliminated. True positive, true negative, false positive, and false negative detections were counted. A keypoint was defined as a correct detection by the network (true positive detection) if there was the corresponding ground truth keypoint in the same image, regardless of its location in the image. For true positive cases, the Euclidean distance between the predicted and ground truth position was calculated as the error of position estimation. The error value represented the normalized value with respect to the length of the monkey’s bounding box due to variations in the size of the monkey in the images. To check the accuracy of the predicted pose, the root-mean-square error (RMSE) was also calculated with all keypoints in each image (Mathis et al.2018). To evaluate the error values of the keypoint position predictions, we investigated human variability by calculating the errors between the keypoint positions annotated by two humans. Finally, among the true positive cases, numbers of limb keypoints misattributed as the homologous keypoint on another limb (e.g., left wrist misattributed as right wrist, left ankle, or right ankle) are also counted. Specifically, *i*-th keypoint were defined as being misattributed to a homologous *j*-th keypoint on another limb, if the keypoint satisfies both of the following two conditions: 1) the normalized position error of the *i*-th keypoint was > 20%; 2) the ground truth positions of *j*-th keypoint was closest to the predicted position of *i*-th keypoint among the ground truth positions of homologous keypoints.

## 3 Results

In total, the present data set contains keypoints and instance segmentation of 16,393 monkeys in 13,083 pictures. Each picture captures 1 to 5 monkeys; 10,630 pictures with a single monkey and 2453 pictures with multiple monkeys (Figure 1).

**Figure 1.**
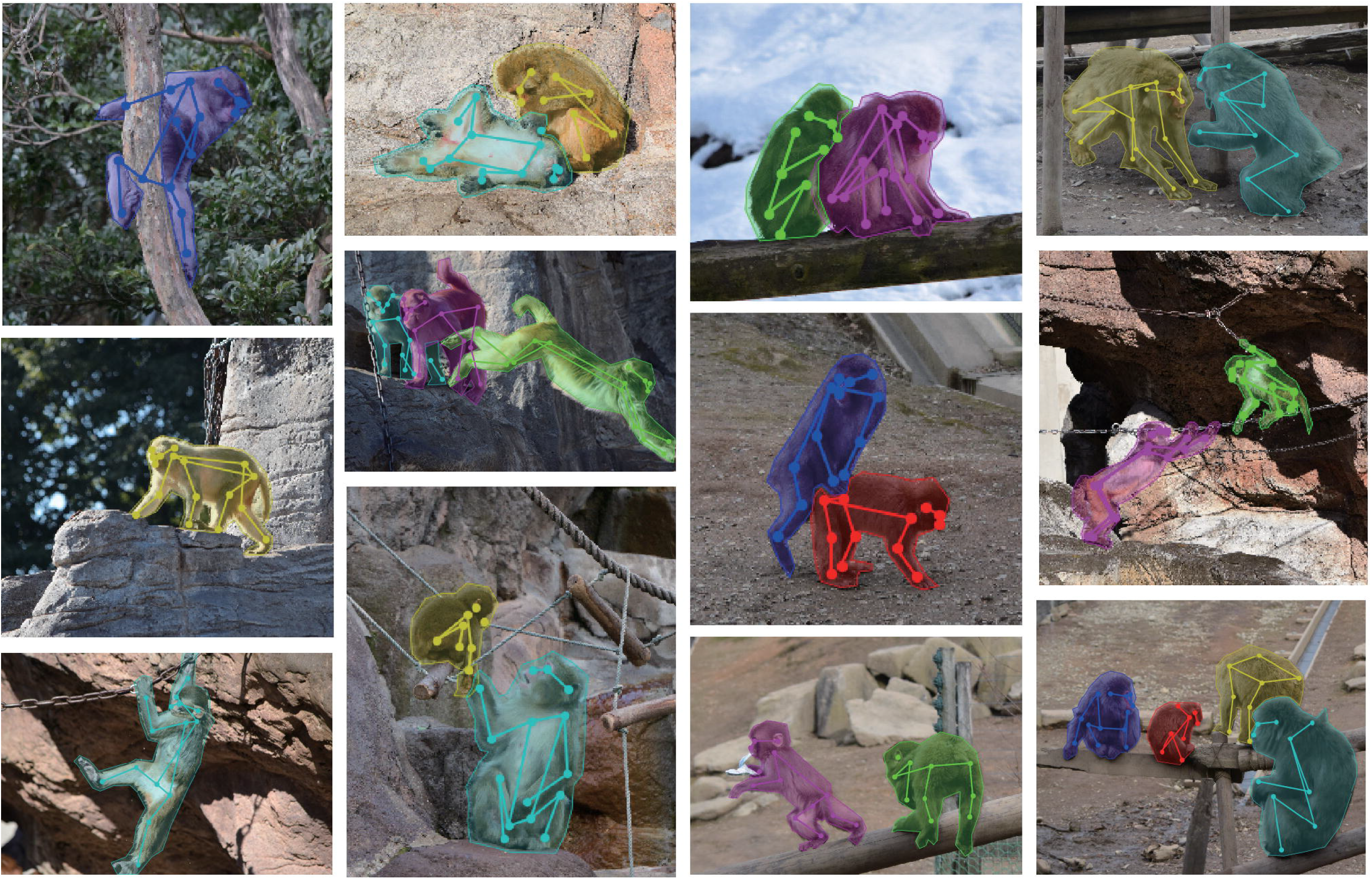
Examples of pictures and labels in the present dataset

To validate the dataset, we trained an artificial network with 14,697 single monkey images in the dataset using the DeepLabCut algorithm (Mathis et al.2018). The performance of the keypoint prediction of the trained network was evaluated on 779 test images unseen during training. Figure 2 shows examples of the keypoint predictions (also see Supplementary Video 1 for keypoint prediction in videos). Among 779 images, 24 images had keypoint(s) detected outside the target monkey. Most of them (17 images) were due to imperfect masks of the other monkeys in the picture (Supplementary Figure 1). The ‘out of monkey’ cases were removed from the analysis.

**Figure 2.**
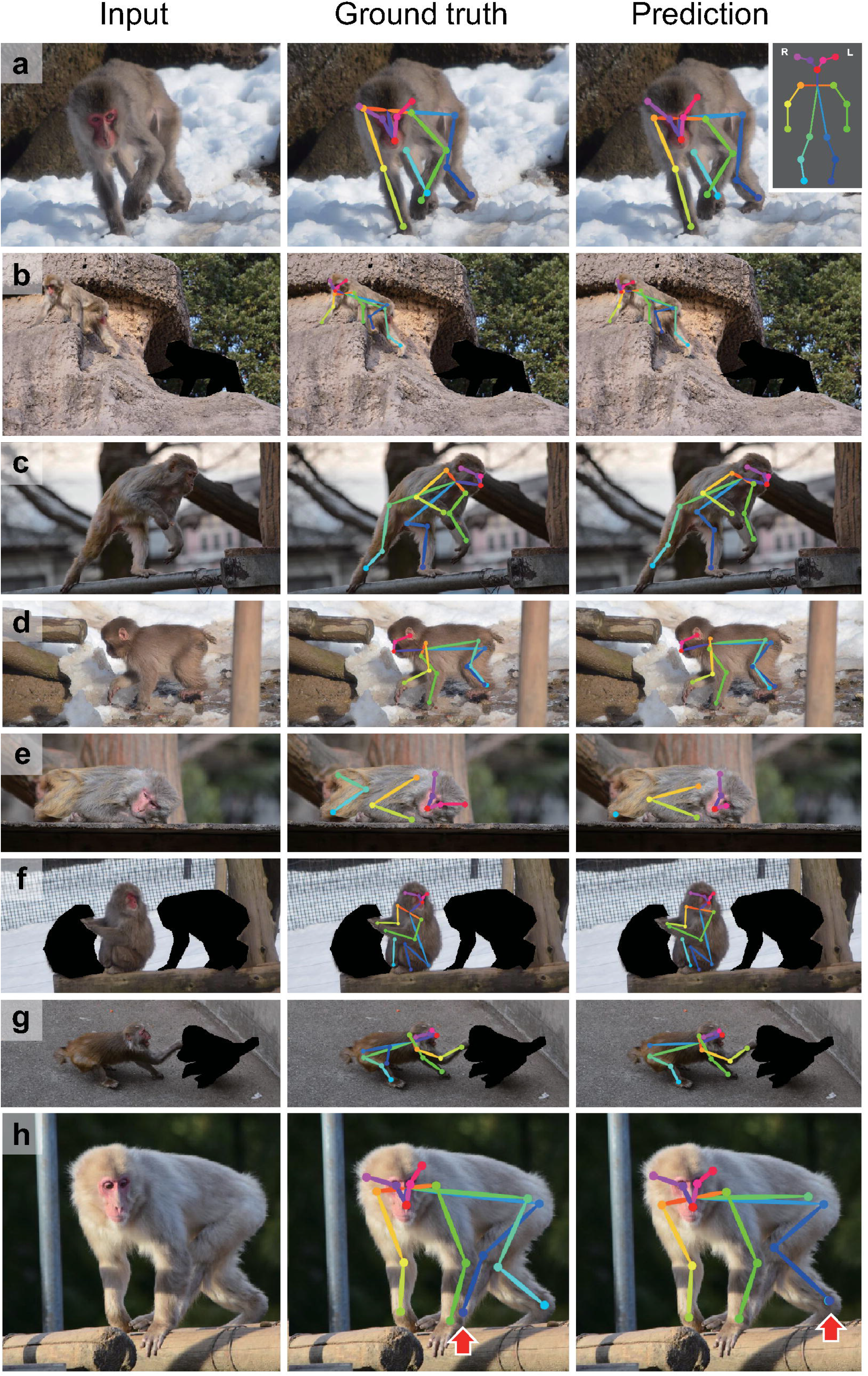
Examples of test image predictions. Test images (left), the ground truth keypoint positions (center) and the position predicted by the artificial neural network trained with the present dataset using the DeepLabCut algorithm (right; a-h). The inset (top right corner) shows color codes of the keypoints. Red arrows in h indicate a misattribution error.

We investigated the performance of keypoint detection (judging whether a keypoint exists anywhere in the picture or not) of the trained network (Supplementary Table 1). Both precision and recall of the keypoint detection were approximately 90% in most of the keypoints, suggesting good detection performance. To further investigate the accuracy of the detected keypoints, the error of predicted position was calculated for each keypoint (Figure 3, gray bar). The prediction’s RMSE values (6.02 ± 0.18%; mean ± s.e.m) were comparable to those between the positions manually labeled by two different people (5.74 ± 0.16%; p = 0.250, student’s t-test), suggesting that the trained network’s performance in the keypoint position estimation was close to the human level. The effect of the label refinement by researchers was also examined. The error values for the dataset before the refinement were calculated as previously mentioned. The analyses revealed that the averaged RMSE values after the refinement (6.02 ± 0.18%) were significantly smaller than the one before the refinement (7.83 ± 0.23%; p = 9.13×10^−10^, Student’s t-test; see Supplementary Figure 2 for the error value of each keypoint). The result suggests that the network trained with the dataset refined by the researchers predicted the keypoint more consistently.

**Table 1.** The number of pictures and monkeys in the present dataset from each source.

| Source | Monkey Species | No. of Pictures | No. of Monkeys |
| --- | --- | --- | --- |
| Toyama Municipal Family Park Zoo | Japanese Macaque | 3784 | 4952 |
| Itozu no Mori Zoological Park | Japanese Macaque | 1312 | 1622 |
| Primate Research Institute | Japanese Macaque | 1641 | 2131 |
| Inokashira Park Zoo | Rhesus Macaque | 2747 | 3203 |
| Tobu Zoo | Rhesus Macaque | 2461 | 2755 |
| Google Open Images | Various | 1138 | 1730 |
| Total |  | 13083 | 16393 |

**Figure 3.**
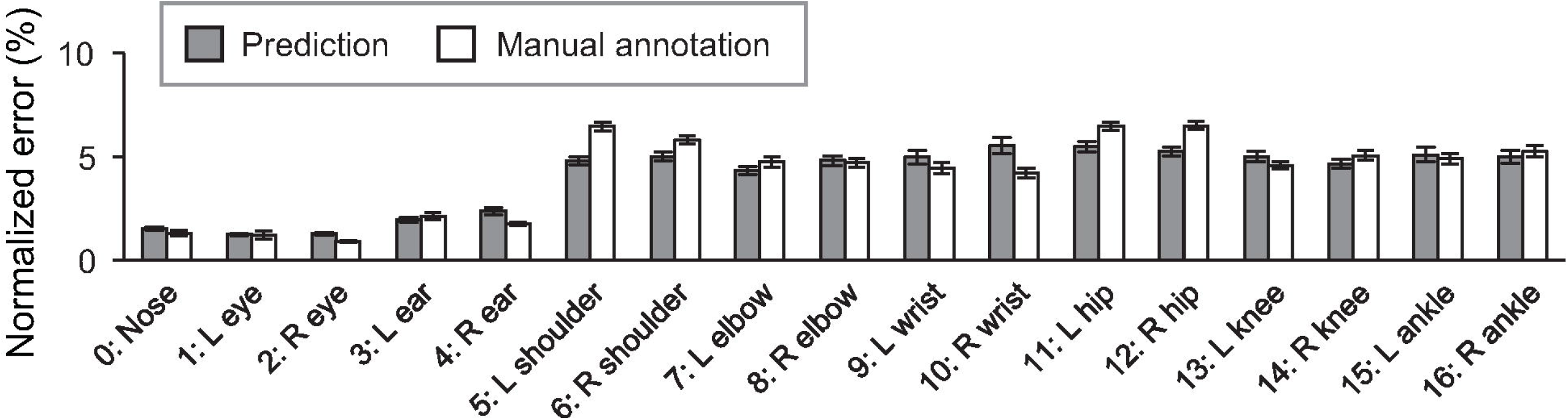
Averaged error of predicted (gray) and manual labeled (white) positions of each keypoint comparing with the ground truth positions. Error bars represent standard error of the mean (s.e.m.)

In some cases, we observed that the predicted positions of monkey’s keypoints on a limb were located on homologous keypoints on another limb (Figure 2h, see also Supplementary Video 1).

We then quantified the frequency of such misattribution errors (Table 2). The misattribution errors were relatively frequent in the distal keypoints (elbow, knee, wrist, and ankle), especially on the hind limbs. The total number of images having at least one misattribution error was 114 (15%). The result shows that there is still room for improvement, although the RMSE indicates human-level performance.

**Table 2.** Number of the misattribution errors. L-R incorrect referring to left or right predicted keypoint was incorrect; F-H switch, forelimb or hindlimb label was incorrect.

| Keypoint pairs | Correct | L-R incorrect | F-H incorrect | L-R and F-H incorrect | Total |
| --- | --- | --- | --- | --- | --- |
| Shoulder | 1225 | 5 | 1 | 3 | 9 |
| Hip | 1062 | 7 | 2 | 3 | 12 |
| Elbow | 1134 | 11 | 5 | 4 | 20 |
| Knee | 1066 | 45 | 11 | 3 | 59 |
| Wrist | 1049 | 14 | 7 | 6 | 27 |
| Ankle | 1045 | 29 | 11 | 5 | 45 |

## 4 Discussions

In this study, we created a novel large dataset of labeled keypoints of macaque monkeys (Figure 1, Table 1). The keypoint estimation performance of the neural network trained with the dataset was close to that of human level (Figure 2, Figure 3, and Supplementary Video 1), demonstrating the usefulness of the present dataset. We also found a significant improvement of the network prediction after the label refinement by researchers using macaques (Supplementary Figure 2), suggesting that the refinement successfully enhanced the quality of the dataset. Although we tested only single monkey images due to the limitation of the algorithm, the present dataset should be useful to train/test the network for multi-animal motion capture. The label formats in the present dataset are compatible with those used in the COCO dataset for humans (Lin et al., 2014), allowing users to try a direct application of algorithms developed for human motion capture. A recent study also proposed a similarly sized labeled dataset of rhesus monkeys (Bala et al., 2020). In the study, they captured freely moving monkeys in a 2.5 m cubic cage with 62 cameras surrounding the cage. The multi-camera system allows to reconstruct 3D pose after manually labeling images simultaneously captured from 3-4 views. Interestingly, the reconstructed 3D pose is projected to the other around 60 views and enables automatically labeling the images from all the views. This cross-view data augmentation allowed them to get labels of around 200,000 monkey images with 33,192 images labeled manually. The critical difference between the two datasets is that pictures in their dataset were taken in a single laboratory environment, our dataset consists of pictures taken in many different naturalistic environments. Thanks to the `in-the-wild’ aspect of the collected pictures, the present data set has rich variations in pose, body shape, lighting, and background in naturalistic contexts. The rich variation will help to train and test artificial neural networks with high generalizability (Mathis et al., 2019). Thus, the two datasets will compensate each other to train or test better neural networks in future studies. As the dataset formats (i.e., which keypoints are labeled) were slightly different among the two datasets, some additional efforts are necessary to combine or compare these two datasets directly.

To understand how the brain generates our behavior, analyzing naturalistic behaviors is crucial. The brain evolved from natural behaviors, and various behaviors, such as complex social behaviors, can be observed only in the natural situations (Datta et al., 2019; Mathis et al., 2020). The high-resolution spatiotemporal data obtained with the markerless motion capture will also aid in understanding brain dynamics underlying the behavior (Berger et al., 2020). Specific posture and motion are informative for studying animals’ emotions and intension (Nakamura et al., 2016), and the motor functions (Berger et al., 2020). Furthermore, the automatic and long-term analyses of naturalistic behavior from a large number of subjects permit new data-driven approaches to find unusual behaviors, personalities and, underlying genetic and neural mechanisms (Vogelstein et al., 2014; De Chaumont et al., 2018). For instance, the recently discovered autistic traits exhibited by macaque monkeys (Yoshida et al., 2016) was identified by such a behavioral observation. Thus, the markerless motion capture for macaque monkeys developed based on the present dataset will be of great use for many neuroscience studies.

The performance evaluation of the network trained with the present dataset revealed that there is still room for improvement regarding the misattribution of the limb keypoints (Figure 2h, Table 2), although the RMSE indicates the human-level performance (Figure 3). The DeepLabCut algorithm (Mathis et al., 2018) used in the present evaluation does not explicitly utilize the prior knowledge about the animal’s body, whereas the other algorithms were suggested to use the connection between keypoints (Insafutdinov et al., 2016; Cao et al., 2017) or 3D shape of the subject (Biggs et al., 2018; Zuffi et al., 2019). Such utilization of the prior knowledge may help to improve the estimation. However, even the state-of-the-art human motion capture algorithms also have difficulties in analyzing the pictures with severe occlusion or crowded people (Mathis et al., 2020). Due to severe occlusions more frequently being observed in naturalistic behaviors in monkeys than in humans, better algorithms may be required in the future. An alternative approach for the improvement will be enriching the dataset itself. Although we tried to capture many different poses in various contexts, the sampling was biased to the frequently observed poses. Adding data selectively for the rarely observed poses may improve the performance of the trained network. Combining with the other monkey datasets made for laboratory environments (Berger et al., 2020; Bala et al., 2020) or transfer learning of the network trained with the human dataset (Sanakoyeu et al., 2020) are also interesting approaches. Nevertheless, in practice, the performance of the network shown in the present study may be sufficient for many applications, after appropriate temporal filtering of the motion data (Berman et al., 2014; Nath et al., 2019) and additional training with the labels made on the pictures in the target experiment (Mathis et al., 2019).

The keypoint position estimation in a 2D video is not the aim of the behavior analysis. In addition, the researchers would need to reconstruct the 3D pose and motion of the animals (Nath et al., 2019; Bala et al., 2020) and label the behaviors that the animals are exhibiting (Datta et al., 2019). The post-processing methods for converting the high-dimensional motion data into meaningful and interpretable behavioral events and parameters of a single animal or interacting animals are still under active developments (Berman et al., 2014; Datta et al., 2019; Dviwedi et al., 2020). The present dataset will permit simple access to motion data of macaques in various environments. Furthermore, this could accelerate the development of post-processing method by accumulating the motion data associated with various natural behaviors.

## 5 Conclusion

We created a novel large open dataset of keypoint labels of macaques in naturalistic scenes. The dataset will be instrumental to train/test the neural networks for markerless motion capture of the macaques and developments of the algorithms for the networks, contributing to the establishment of an innovative platform of behavior analysis for non-human primates for neuroscience and medicine, as well as the other fields using macaques (Carlsson et al., 2004).

## 6 Conflict of Interest

The authors declare that the research was conducted in the absence of any commercial or financial relationships that could be construed as a potential conflict of interest.

## 7 Author Contributions

RL, TS, JM, KI, YG, and HNishij designed this research. JM, KI, TS, RL, and MT created the dataset. RL, SN, JM, TS and HNishim evaluated the performance of the neural network trained with the dataset. All the authors discussed the results and commented on the manuscript, read and approved the final manuscript.

## 8 Funding

This work was supported by the Cooperative Research Program of Primate Research Institute, Kyoto University, the Grant-in-Aid for Scientific Research from Japan Society for the Promotion of Science (No. 16H06534 and 19H04984), and the grant of Joint Research by the National Institutes of Natural Sciences (NINS) (NINS Program No. 01111901).

## 9 Acknowledgments

We would like to thank Yu Yang, Yang Meng, Kei Kimura, Yukiko Otsuka, Andi Zheng, Jungmin Oh, Gaoge Yan, and Yuki Kinoshita for manually labeling and refining the keypoints.

## Data Availability Statement

The dataset for this study will be released when the manuscript is accepted.

## Notes

### Competing Interest Statement

The authors have declared no competing interest.

### Summary of Updates

Supplemental Files updated

