## Supplemental Figure 1 for "MacaquePose: A novel ‘in the wild’ macaque monkey pose dataset for markerless motion capture"

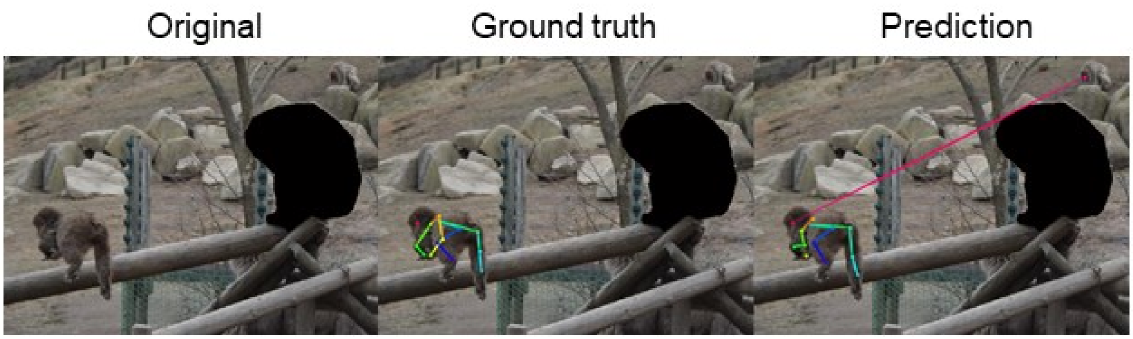


**Supplementary Figure 1.** An example of 'out of monkey' keypoint detection error. Keypoint detection error due to the imperfect masking of the other monkeys.
