## Supplemental Figure 2 for "MacaquePose: A novel ‘in the wild’ macaque monkey pose dataset for markerless motion capture"

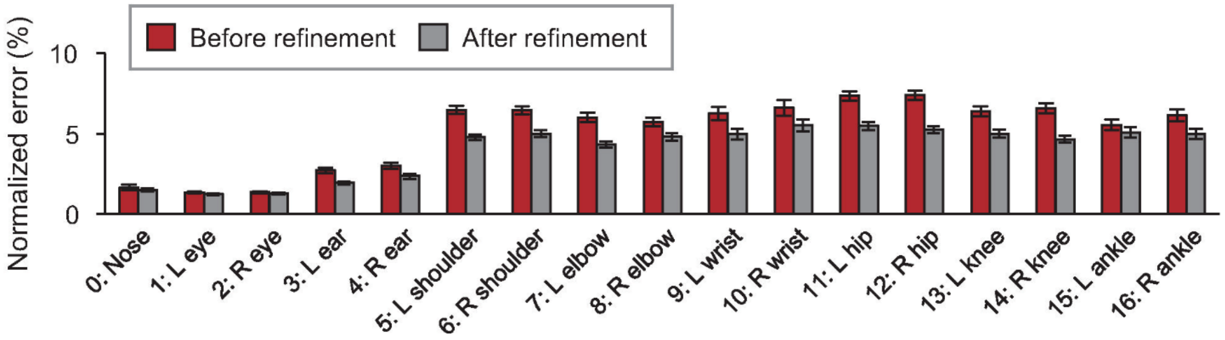


**Supplementary Figure 2.** Averaged error observed before and after the researchers' refinement. Averaged error of prediction by the network trained with the dataset before (red) and after (gray) the refinement by researchers. Error bars represent standard error of the mean.
