## Supplemental Table 1 for "MacaquePose: A novel ‘in the wild’ macaque monkey pose dataset for markerless motion capture"

**Supplementary Table 1.** Summary of keypoint detections. Values inside cell are number of test images, except specified in %. Precision = TP/(TP + FP); Recall = TP/(TP + FN)

| Keypoints | True Positive, TP | True Negative, TN | False Positive, FP | False Negative, FN | Precision in % | Recall in % |
|---|---|---|---|---|---|---|
| Nose | 543 | 171 | 62 | 9 | 89 | 98 |
| Left Eye | 393 | 279 | 69 | 14 | 85 | 97 |
| Right Eye | 403 | 272 | 72 | 8 | 85 | 98 |
| Left Ear | 433 | 222 | 78 | 22 | 85 | 95 |
| Right Ear | 423 | 199 | 106 | 27 | 80 | 94 |
| Left Shoulder | 613 | 33 | 56 | 53 | 92 | 92 |
| Right Shoulder | 621 | 32 | 69 | 33 | 90 | 95 |
| Left Elbow | 565 | 90 | 54 | 46 | 91 | 92 |
| Right Elbow | 589 | 78 | 47 | 41 | 93 | 93 |
| Left Wrist | 553 | 100 | 50 | 52 | 92 | 91 |
| Right Wrist | 572 | 88 | 52 | 43 | 92 | 93 |
| Left Hip | 553 | 72 | 73 | 57 | 88 | 91 |
| Right Hip | 521 | 92 | 61 | 81 | 90 | 87 |
| Left Knee | 543 | 93 | 64 | 55 | 89 | 91 |
| Right Knee | 533 | 108 | 58 | 56 | 90 | 90 |
| Left Ankle | 540 | 87 | 71 | 57 | 88 | 90 |
| Right Ankle | 550 | 101 | 60 | 44 | 90 | 93 |
